# Horizontal gene transfer drives the evolution of dependencies in bacteria

**DOI:** 10.1101/836403

**Authors:** Akshit Goyal

## Abstract

Many naturally-occurring bacteria lead a lifestyle of metabolic dependency, i.e., they depend on others for crucial resources. We do not understand what factors drive bacteria towards this lifestyle, and how. Here, we systematically show that horizontal gene transfer (HGT) plays a crucial role in the evolution of dependencies in bacteria. Across 835 bacterial species, we map gene gain-loss dynamics on a deep evolutionary tree, and assess the impact of HGT and gene loss on bacterial metabolic networks. Our analyses suggest that genes acquired by HGT can affect which genes are later lost. Dependency evolution is contingent on earlier HGT because of two reasons. First, we find that HGT typically adds new catabolic routes to bacterial metabolic networks. This increases the chance of new metabolic interactions between bacteria, which is a prerequisite for dependency evolution. Second, we show that gaining new routes can promote the loss of specific ancestral routes (a mechanism we call “coupled gains and losses”, CGLs). Phylogenetic patterns indicate that both types of dependencies — those mediated by CGLs and those purely by gene loss — are equally likely. Our results highlight HGT as an important driver of metabolic dependency evolution in bacteria.

---

Naturally-occurring bacteria lead one of two metabolic lifestyles: autonomy or dependency [1–4]. While autonomy reflects complete self-sufficiency in converting nutrients to biomass, bacteria with dependencies often require crucial metabolites from others [5–9]. These metabolites are secreted by neighbouring community members. Since such dependencies are common in bacterial communities, it is instructive to ask: what processes and factors affect their evolution; in other words, what drives the switch from metabolic autonomy to dependency?

To answer this question, recent studies have put forth the Black Queen hypothesis, which states that dependencies evolve through adaptive gene loss [10–12]. Individuals lose costly, dispensable genes in leaky environments, trading autonomy for better growth (or fitness). As both experiments and models show, this is feasible — administering the loss of even a few specific biosynthetic genes in bacteria repeatedly leads to strong metabolic dependencies [13–16]. This also explains how endosymbionts undergo severe genome reduction [2]. These bacteria lack many biosynthetic pathways, and instead depend on their hosts for the required biomass components.

However, many extant free-living bacterial species are also metabolically dependent, despite the “free-living” label [17–21]. These species do not have merely reduced genomes, i.e., they do not differ from their ancestors only by gene losses, as expected under the Black Queen hypothesis [22–24]. Over time, they have also gained many genes, primarily by horizontal gene transfer (HGT) [25–29]. For these often-dependent bacteria, we ask: could gene gains also have contributed to which dependencies we observe today? Specifically, during dependency evolution, can which genes are gained influence which genes will later be lost?

Here we explore the role of horizontal gene transfer in the evolution of metabolic dependencies in bacteria. Specifically, we measure the potential of HGT to drive dependency evolution by affecting the likelihood of subsequent gene loss events. To do this, we reconstructed the evolutionary history of 835 phylogenetically diverse, non-endosymbiont bacterial species. By inferring ancestral gene content, we mapped gene gains and losses along a large, deep-branching phylogeny, and assessed their impact on bacterial metabolic capabilities. Our analyses suggest that horizontally transferred genes can indeed affect which genes are later lost, and which dependencies emerge as a result. We have two lines of evidence to support this. First, we find that gene gains add new catabolic routes to bacterial metabolic networks. These gained catabolic routes increase the chance of new metabolic interactions between bacteria, a prerequisite for dependency evolution. Next, we show how these new routes can promote the loss of pre-existing ancestral routes (a process we call “coupled gains and losses”, CGLs). We find that phylogenetic patterns indicate that both processes — CGLs and pure gene loss — are equally likely to lead to dependencies. Collectively, these results highlight horizontal gene transfer as an important driver of metabolic dependency evolution in bacteria.

## RESULTS

### Horizontal gene transfer (HGT) adds new catabolic routes to bacteria

In a metabolic dependency, a donor organism secretes metabolites, which are in turn required by an acceptor organism. The secreted metabolites cannot be produced by the acceptor organism itself, but are still necessary for survival and growth. We sought to understand how horizontal gene transfer, if at all, impacts the emergence of new dependencies. We hypothesized that newly acquired genes (through HGT) lead to newer metabolic interactions. This could occur if gained genes allowed an acceptor organism to transport and break down previously unusable metabolites in its surroundings. We thus first asked: does horizontal gene transfer enhance the ability of a bacterial genome to utilize metabolites secreted by surrounding donors? In other words, does it add new catabolic routes?

To answer this, we used a two-pronged approach, combining bacterial metabolism and phylogeny. We first acquired a list of 1,031 bacterial species with complete genomes, whose metabolic data were available in the KEGG GENOME database. We explicitly removed from this list: (1) endosymbionts, due to their exotic metabolic lifestyles and genomes, and (2) closely related genomes, to avoid phylogenetic bias (see Methods). This left us with the 835 species genomes we used for all our subsequent analyses (supplementary table 1). For each genome, we extracted all metabolic genes present in at least one species, corresponding to a total of 3,022 unique genes.

Using these genomes, we inferred each species’ metabolic capabilities. For this, we reconstructed representative metabolic networks, one for each species, using gene presence-absence data (figure 1a). Here, we mapped each gene to specific chemical reactions using the KEGG REACTION database (see Methods). To identify gene gain and loss events during the evolution of these species, we inferred the most likely genetic makeup of their ancestors. For this, we first established evolutionary relationships using a well-known bacterial phylogenetic tree, and then applied ancestral reconstruction methods, to infer which of the 3,022 metabolic genes were likely present in each ancestor (see Methods). With the presence-absence profiles of ancestors and descendants on each phylogenetic branch, we could infer which genes were gained and lost along them (figure 1b).

**FIG. 1.**
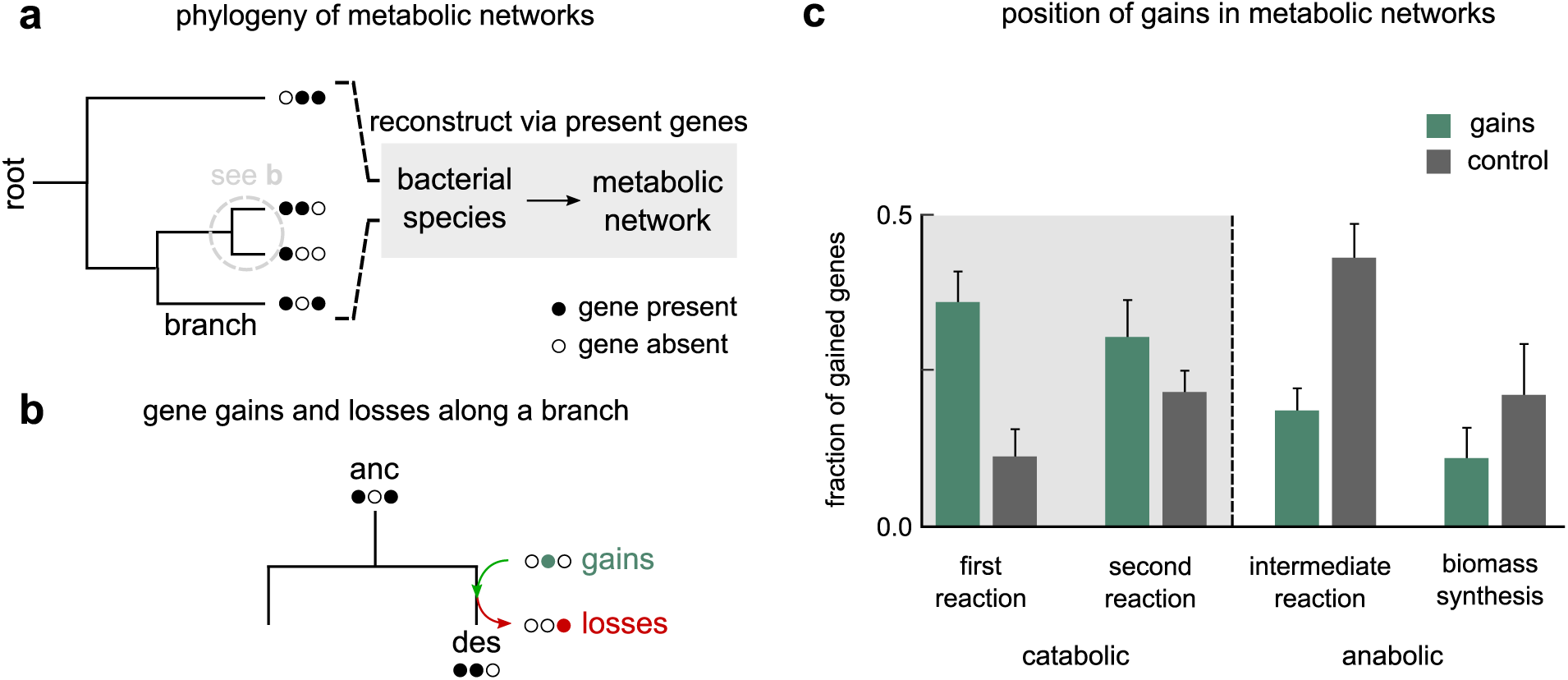
Horizontal gene transfer adds new catabolic routes to bacteria. **a**, Schematic representation of our two-pronged approach: of combining phylogeny and bacterial metabolism. We used a well-known phylogenetic tree to infer the evolutionary relationships between the 835 bacterial species used in our analysis. For each extant species (shown on the tips of the tree), we used gene presence-absence data for 3,022 metabolic genes from the KEGG metabolic database. Filled circles indicate gene presence; empty indicate absence. **b**, We inferred the gene presence-absence states of all internal nodes of the tree along each branch of the full tree (grey dashed circle in **a**); each branch connected an ancestor (*anc*) to a descendant (*des*). Along each branch, we inferred which genes were gained (green) and lost (red). **c**, Bar plot showing the position of gained genes in bacterial metabolic networks. We split metabolic genes into catabolic (first or second reactions in a metabolic route) and anabolic (intermediate or biomass-synthesis reactions) based on the the chemical reactions they map to. Each green bar represents the average fraction of gained (horizontally transferred) genes at that position. Each black bar represents controls, i.e., the expected average fraction of gains at that position, given a random set of gene gains. Error bars show the standard error, indicating the extent of variation across 1,669 phylogenetic branches.

We first tested whether HGT can expand the set of externally available metabolites that a metabolic network can catabolize. For this, we studied which position horizontally transferred genes typically occupied in each metabolic network they were gained in. Specifically, we were interested in whether the gained genes were in catabolic or anabolic parts of a metabolic network. We studied this across all phylogenetic branches. On each branch, we asked which position in the descendant’s metabolic network each gained gene occupied. Each position corresponded to metabolic reaction order, from catabolic to anabolic, as follows: first, second, intermediate or biomass-synthesis reactions (see Methods). If HGT was indeed likely to add new catabolic routes, we would expect gained genes to be concentrated in the catabolic parts of the network, i.e., first and second reactions. As controls, we measured the positions of randomly chosen genes in the same metabolic networks.

Along each branch, we measured the fraction of gained genes corresponding to each network position. We then calculated, across all branches, the average fraction of gains found to occupy each position. Consistent with our hypothesis, we found that horizontally acquired genes are more likely to be part of catabolic routes (mean; 69%catabolic versus 31% anabolic; figure 1c, green bars); this is much more than expected by chance (in controls, we found mean 34% catabolic versus 66% anabolic; figure 1b, black bars). This suggests that HGT can expand the number of external metabolites that bacteria can catabolize.

### HGT-enabled catabolic routes increase the likelihood of metabolic interactions

We next tested the possibility that newly acquired catabolic routes promote new metabolic interactions. For this, new routes should help break down the metabolic byproducts of other bacteria more often than nutrients available in the environment. To test this, we first curated a list of common external metabolites, and classified them as byproducts or nutrients based on their presence on the exterior or interior of microbial metabolic networks (see supplementary table 2 for the full list; see Methods for a detailed procedure). In a metabolic interaction, the received metabolite should eventually help produce biomass components (such as pyruvate, ribose-5-phosphate, and alanine). For each ancestor-descendant pair, we analyzed how many such biosynthetic routes were added to the descendant’s metabolic network, when compared with its ancestral network (see Methods). We separately counted routes using byproducts as their starting point, from routes using nutrients.

We found that, on average, new biosynthetic routes, enabled by HGT, are more likely to be byproduct-driven than nutrient-driven (median number of routes, 56 and 51, respectively; *P* < 10^−3^; distributions compared via a Kolmogorov-Smirnov test; see figure 2a). Moreover, these new routes could often be metabolically more efficient than their ancestral counterparts (see figure 1d), i.e., they often had shorter path lengths (49%; figure 2b, left) and yielded more energy (ATP; 58%; figure 2c, right) than the corresponding routes in their ancestors (see Methods). This suggests that newly acquired routes can indeed enable new metabolic interactions with other donor bacteria. In fact, some of these interactions can also have adaptive significance, which can encourage the evolution of metabolic dependencies.

**FIG. 2.**
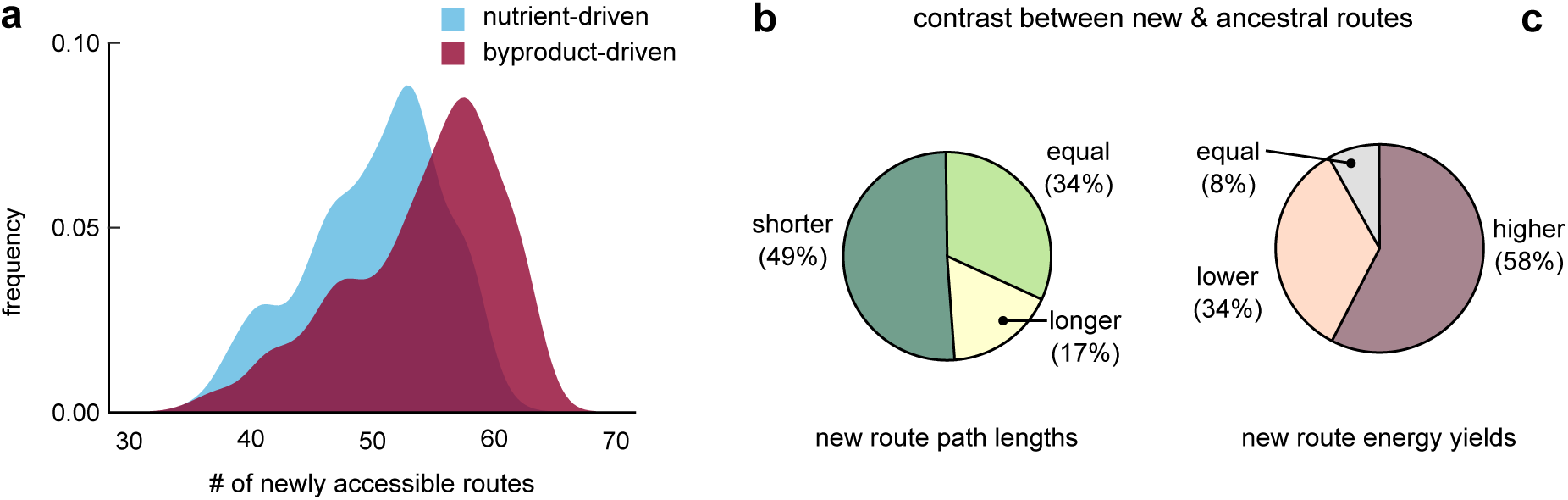
HGT-enabled catabolic routes increase the likelihood of metabolic interactions. **a**, Distribution of the number of newly accessible catabolic routes (routes gained along each phylogenetic branch) across all 1,669 branches. The number of routes starting from nutrients are shown in blue, and those starting from byproducts are in red. New byproduct-driven routes would increase the chance of metabolic interactions with other bacteria (via their byproducts). We find that new catabolic routes are more likely to be byproduct-driven (median 56, versus 51 for nutrient-driven routes; *P* < 10^−3^; Kolmogorov-Smirnov test). **b–c**, Pie charts comparing new routes with their corresponding ancestral routes on each of the 1,669 branches. We compare new and ancestral routes based on their **b**, path length (i.e., is the shortest new path shorter, longer or the same length as the shortest ancestral path), and **c**, energy yield (i.e., does the most energy-yielding new path have a higher, lower or equal yield than the best ancestral path).

### HGT can affect dependency evolution via coupled gains and losses of genes

Given that newly acquired routes have similar — and sometimes better — path lengths and energy yields, we wondered if their acquisition could promote the loss of corresponding ancestral routes. We thus hypothesized the following mechanism through which, contingent on earlier HGT, metabolic dependencies could emerge by subsequent gene loss.

Consider the example illustrated in figure 3a, with an environment that consists of a nutrient (*nut*, blue circle), and a byproduct (*byp*, purple triangle) secreted by a donor (not shown). Consider a specific acceptor organism, that requires the biomass component (*bmc*, yellow square) either directly or indirectly, to survive. We follow the modification of this acceptor’s metabolic network, from ancestor to descendant, in three steps. First, the ancestor uses a specific metabolic pathway (labelled “ancestral route”) to convert the available nutrient to the essential biomass component. Second, it gains a catabolic route (labelled “gained route”) that uses the byproduct, *byp*, to produce *bmc*. Third, after such a gain, the acceptor loses the ancestral route to *bmc*. This is allowed because *bmc* — crucial for survival — can still be produced through a coupled (or alternate), byproduct-utilizing route. Once lost, however, the acceptor will become obligately dependent on its neighbours to receive the byproduct.

**FIG. 3.**
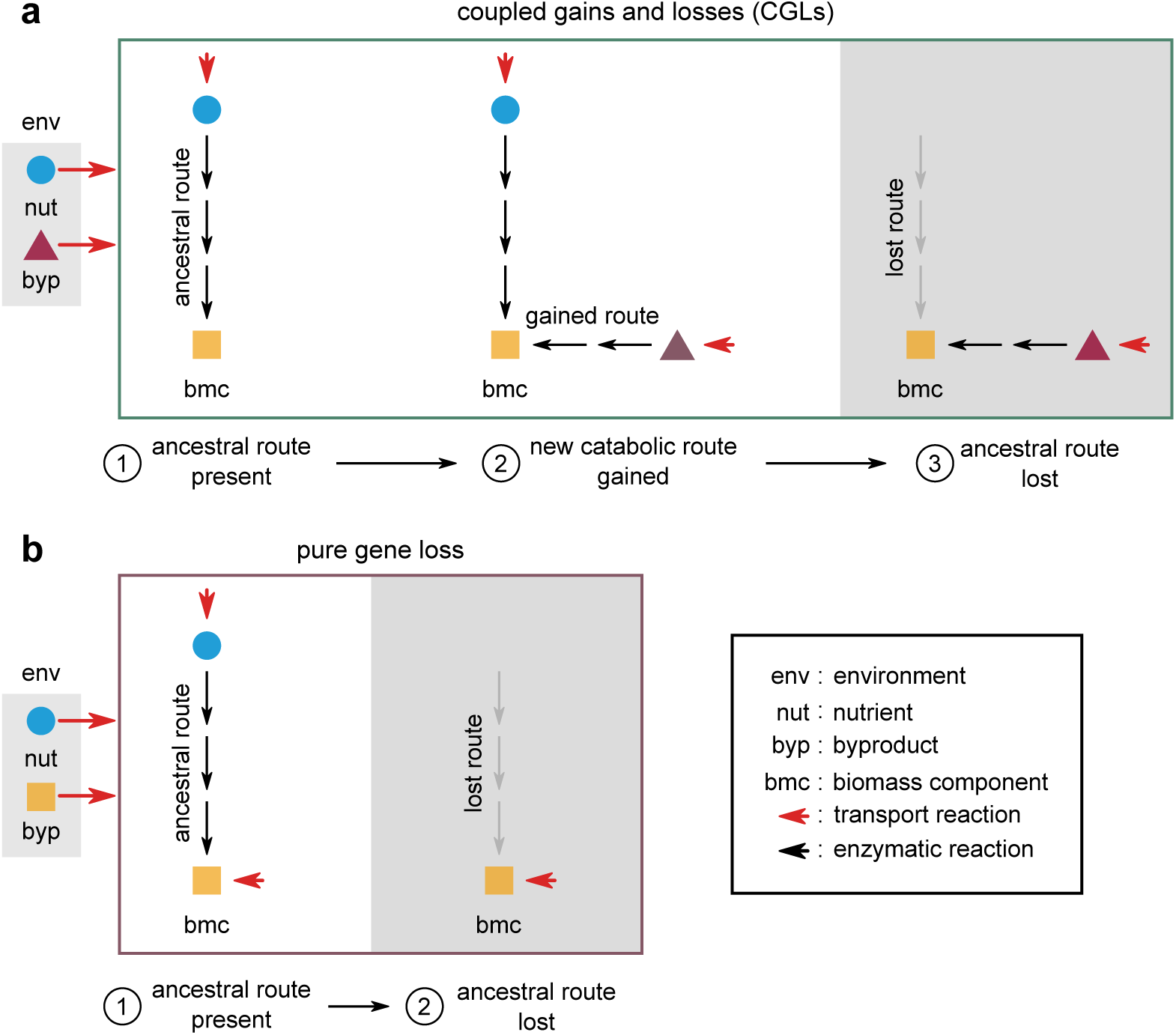
HGT can affect dependency evolution via coupled gains and losses of genes. **a**, Schematic illustration of coupled gains and losses (CGLs), a new, alternate mechanism for dependency evolution driven by HGT. The grey box on the left indicates the environment, with a nutrient (*nut*, blue circle) and a byproduct (*byp*, purple triangle). Red arrows indicate the secretion and import of metabolites by bacteria. The three steps in the frame show how a bacterial species can evolve a dependency on another species that donates *byp* in the environment; by step 3, the acceptor species eventually depends on *byp*. Each step follows the modification of a part of the acceptor’s metabolic network. At step 1, the acceptor uses an ancestral route (black arrows) to convert *nut* in the environment to a key biomass component (*bmc*, yellow square). At step 2, it can alternatively use a newly gained route to convert *byp* to the same *bmc*. The ancestral and gained routes are coupled to each other, because they both produce *bmc*, which is crucial for survival and growth. At step 3, the acceptor loses the ancestral route (grey arrows), but can still produce *bmc* through the gained route. It thus becomes dependent on donors of *byp* for survival. **b**, Schematic illustration of dependency evolution via pure gene loss, not driven by HGT. The environment now has *bmc* available as a byproduct, instead of *byp* (grey box on the left). In this mechanism, step 1 is the same as **a**, but unlike **a**, at step 2, the acceptor can lose the ancestral route straight away, without requiring an alternate coupled route. However, this requires a particular environment where *bmc* is available.

Note that here, until the gain of an alternate route, the ancestral route cannot be lost by the acceptor. Moreover, gaining such a route can not only allow the loss of the ancestral route, but also promote it. This is most likely, for instance, when the gained route is more efficient than the pre-existing route (which is often the case; figure 2c). Collectively, we term this process (figure 3a, steps 1 to 3), coupled gains and losses (CGLs). CGLs demonstrate how HGT can crucially affect metabolic dependency evolution.

Contrast CGLs with pure gene loss, which relies on the environmental availability of *bmc* for dependency evolution (figure 3b). Unlike CGLs, these events are unlikely to be affected by, or depend on, HGT. Interestingly, the same genome (or microbial species) can evolve the same dependency via both mechanisms, but their likelihoods are crucially environment-dependent. This is transparent in figure 3a, b, where the primary difference between the two cases is which byproduct is available: in figure 3a, it is *byp* (purple triangle); in figure 3b, it is *bmc* (yellow square). The only other difference is the gain of a route to metabolize *byp*, which is likely in an environment where *byp* is available as a byproduct.

### Metabolic dependencies are equally likely to emerge via CGLs and pure gene loss

Given that both coupled gains and losses (CGLs) and pure gene loss are possible mechanisms for metabolic dependency evolution, we asked which of them was more likely to cause the metabolic dependencies observed in extant bacteria. To help answer this, we looked for two distinct, but related phylogenetic signatures.

First, we measured what fraction of ancestor-descendant transitions (each represented by a phylogenetic branch) displayed gain-loss patterns consistent with CGLs; we compared this with the fraction of transitions consistent with pure gene loss. Here, for CGL-consistent transitions, we asked whether a species gained a catabolic route for at least one biomass component which it also lost an alternate route for, during the ancestor-descendant transition (i.e., along a phylogenetic branch, how often did we detect any CGL events, like in figure 3a; see Methods). For transitions consistent with pure gene loss, we similarly asked how often we observed a species losing a pre-existing route, without gaining an alternate catabolic route for the same biomass component (i.e., along a phylogenetic branch, how often did we detect any pure gene loss events, like in figure 3b; see Methods). To control for the likelihood of both events occurring by chance, we also calculated the expected fractions of CGL and pure gene loss events, by repeating our measurements on simulated datasets (see Methods). In each simulated dataset, we randomized which genes were gained and lost along each ancestor-descendant transition, or branch. Here, we ensured that the same number of genes were gained and lost along each branch as those in our observed dataset. We found at least one CGL event on 33% of all branches (figure 4a; green); in comparison, we found pure gene loss events on 24% of all branches (figure 4a; red). Both kinds of events were observed much more likely than expected by chance (figure 3c; greys). This suggested that by merely observing gene gain-loss patterns, we would conclude that CGLs were more likely to lead to dependencies than pure gene loss.

**FIG. 4.**
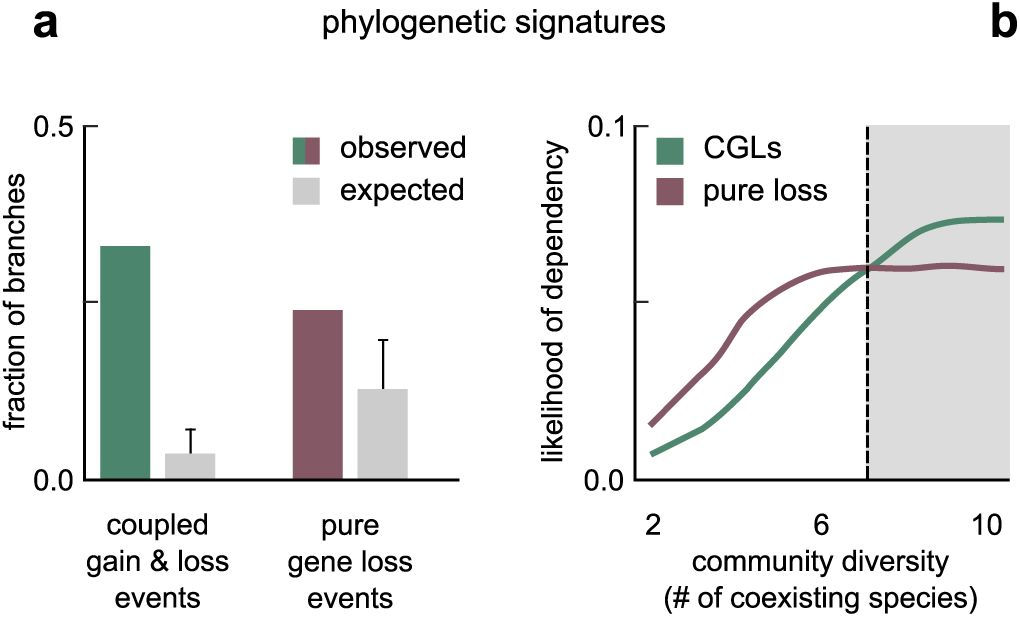
Metabolic dependencies are equally likely to emerge via CGLs and pure gene loss. **a**, Bar plot showing the fraction of 1,669 phylogenetic branches in which we observed gene gain-loss patterns consistent with coupled gains and losses (CGLs; green), and with pure gene loss (red). Each grey bar represents the corresponding controls, i.e., the expected fraction of branches with patterns consistent with CGLs and pure gene loss, given a random set of gene gains and losses. Error bars show the standard error, indicating the level of variation across all branches. **b**, Line plot showing the likelihood of evolving dependency via CGLs (green) and pure gene loss (red) in simulated bacterial communities, as a function of the community diversity. The likelihood of dependency is the average fraction of communities in which the observed gains and losses along a branch led to a CGL-based on pure gene loss-based dependency. Community diversity is the number of coexisting bacterial species in a simulated community. The grey region has ≥ 7 species, where CGLs are more likely than pure gene loss.

Second, we added environmental context to the observed gain-loss patterns, by measuring how the likelihood of metabolic dependency evolution depended on a species’ microbial community. Specifically, we asked how the chance of both events — CGLs and pure gene loss — depended on the number of species in a microbe’s community (hereafter, community diversity). We hypothesized that more diverse communities would have a higher number of available nutrients in the environment, because more species would secrete metabolic byproducts. We expected that increasing community diversity would thus generally favour dependency evolution via both CGLs and pure gene loss; we did not know which of the two would be more favoured. To measure the effect of community diversity on the chance of CGLs and pure gene loss across several environments, we asked how often the observed gains and losses would lead to dependencies because of CGLs and pure gene loss alone across hundreds of thousands of random, simulated microbial communities. We used simulated communities as proxies for environments, due to our lack of knowledge of the actual environments of different species across their evolutionary histories; in doing so, we were estimating the typical chance of dependencies emerging via CGLs and pure gene loss. For these simulations, we curated 1,035 environments, each with a different pair of nutrients present (supplementary table 3; see Methods for details). In each environment, we randomly chose unique sets of bacterial species from the 835 in our study as different communities; we chose 100 unique species sets at each level of diversity, from 2 to 10 (see Methods).

For each phylogenetic branch, and for each environment-community pair (roughly 100,000 per level of community diversity), we measured how often the observed gains and losses along the branch led to a new dependency in the descendant; we measured this separately for CGLs and pure gene loss. Briefly, a dependency was CGL-mediated when the following three conditions were satisfied: (1) the descendant gained an alternate route while also losing a coupled route (similar to figure 3a), (2) the gained route used a metabolic byproduct from the community for biomass production, and (3) the biomass component produced via the gained route was not available as a community byproduct. In contrast, a dependency was pure gene loss-mediated when: (1) the descendant lost a pre-existing route for the production of a biomass component (similar to figure 3b), and (2) that biomass component was directly available as a metabolic byproduct from the community (see Methods).

To illustrate our results, we plotted the fraction of simulations where we detected dependency evolution as a function of community diversity, i.e., we plotted the likelihood of dependencies via CGLs and pure gene loss with increasing community diversity (figure 4b). Consistent with our hypothesis, we found that the likelihood of dependencies via both CGLs and pure gene loss increased with increasing diversity (figure 4b; CGLs in green; pure gene loss in red); the likelihood of both events saturated at high diversity. Strikingly, while pure gene loss was more likely at low diversity (< 7; white region in figure 4b), we found that CGLs were more likely at high diversity (≥ 7; grey region in figure 4b). This was because: while the number of byproducts increase with increasing diversity (supplementary figure 1), byproducts are more likely to be pathway intermediates (favouring CGLs) than biomass components (favouring pure gene loss); see supplementary figure 2. Collectively, our analyses and simulations suggest that CGLs and pure gene loss are equally likely mechanisms for metabolic dependency evolution in bacteria.

## DISCUSSION

To summarize, here we showed that horizontal gene transfer (HGT) can play a significant role in metabolic dependency evolution in bacteria. Specifically, if an alternate metabolic pathway (or “route”) is gained by HGT, it can promote the loss a pre-existing, otherwise indispensable route. Such alternate routes often catabolize metabolic byproducts from coexisting bacteria, thus making bacteria dependent on them. Overall, this is a new mechanism for dependency evolution: coupled gains and losses (CGLs). Phylogenetic evidence suggests that CGLs have occurred much more frequently across bacterial evolutionary history than expected by chance (figure 4a). Further, phylogenetic evidence also suggests that CGLs can often be adaptive, since gained pathways are often shorter and more energy-efficient when compared with pre-existing pathways (figure 2b–c).

As a mechanism for metabolic dependency evolution, CGLs are contrasted with pure gene loss, also called the Black Queen hypothesis. We found that while in communities with low diversity, pure gene loss is the more likely cause of dependencies, in communities with high diversity, CGLs are more likely (figure 4b). Our results thus enrich and supplement the Black Queen hypothesis, by explaining the role of prior gene gains on eventual gene loss.

We believe that our approach can also aid in a more accurate classification of bacterial lifestyles. Conventionally, bacteria are classified as either free-living or symbiotic in biological databases. While this classification suggests that free-living bacteria would often be independent (and symbiotic ones, dependent) these labels can be misleading. For instance, free-living bacteria are often metabolically dependent [17]. In our analyses, we wanted to avoid relying on such a binary classification. We acknowledged that the degree of dependency of a bacterial genome lies along a spectrum, and measured it by inferring which key biomass components each genome could synthesize in various nutrient environments. In this way, our approach is more precise and ecologically relevant.

The mechanism we proposed, CGLs, also makes the following prediction about experimental evolution: when co-evolved in a diverse community, bacteria are more likely to lose biosynthetic pathways that they have alternate pathways for; this is less likely when they are evolved alone. As a corollary, adding alternate pathways to bacteria will promote the loss of pre-existing pathways. Both predictions can be tested via laboratory evolution in a community context.

The framework we used here, combining phylogenetic analyses with metabolic network analyses, can also help quantify the relative contributions of drift and selection to the reduction of bacterial genomes. Progressive genome reduction is often termed “genome streamlining”, and a key question in bacterial genome evolution asks how parallel, or repeatable, streamlining events are. The logic is that more parallel events reflect selection being dominant in genome reduction. We can systematically study these questions within our framework. For instance, we can measure how often we detect the same dependencies evolve along a phylogenetic branch, and quantify how similar the corresponding gene loss events are. Similar, or repeatable, gene loss events would be consistent with selection playing a major role in streamlining: perhaps “weeding out” genes no longer required in certain environments. Dissimilar gene loss events, on the other hand, would suggest that drift dominates. Such analyses are outside the scope of this study, and the subject of future work.

Finally, our analyses focused on changes in metabolic network architecture, but dependency evolution can also occur via changes to gene regulatory networks. In experiments, we observe that both metabolic and regulatory changes are responsible for evolved dependencies [15, 30, 31]. However, we do not understand how to incorporate the effect of regulatory changes on bacterial phenotypes as well as we do the effect of metabolic changes. Future work in this direction can help better understand the role of regulation on metabolic dependency evolution.

## METHODS

### Mapping genomes to metabolic networks

We extracted a list of all 1,031 bacterial species whose complete genomes were available in the Kyoto Encyclopedia of Genes and Genomes (KEGG) GENOME database [32]. We then pruned this list to remove endosymbionts and closely related genomes. To remove endosymbionts, we used a curated list of endosymbiont genomes based on literature surveys [33]. To remove closely related genomes, when multiple genomes were available for a species, we chose one at random. This resulted in 835 genomes or species, which we used for all subsequent analyses (see supplementary table 1 for the full list). To infer which metabolic genes were present and absent in each species, we extracted a list of all the genes in that species which mapped to a corresponding metabolic reaction in the KEGG REACTION database. We found a total of 3,022 unique genes that were present in at least one of the 835 species in our dataset. We assumed that the set of all mapped metabolic reactions per species was its metabolic network (figure 1a).

### Inferring gene gains and losses

To obtain species’ phylogenetic relationships, we mapped our 835 genomes to a well-known phylogenetic tree by matching their GenBank accession numbers [34, 35]. To infer the most likely genetic make-up of each ancestor, i.e., the internal nodes of the phylogenetic tree, we used a previously published ancestral state reconstruction method by Cohen and Pupko [36]. Such a method is commonly used to study the long-term evolutionary history of gene gains and losses on deep phylogenetic trees [25, 27]. The parameters we used while running the method were consistent with previous large-scale studies of bacterial genome evolution [28]; the full set of parameters is available in supplementary table 4. We then calculated which genes were gained and lost along each phylogenetic branch, i.e., between an ancestor and its descendant(s), by comparing their gene presenceabsence profiles. We assumed a gene was gained along a branch if it was absent in the ancestor, but present in the descendant; similarly, we assumed a gene was lost along a branch if it was present in the ancestor, but absent in the descendant (figure 1b). We verified that using an alternate, maximum-likelihood based method to infer gene gains and losses did not significantly affect our results (< 5% mismatch between gain-loss patterns across all 1,699 branches).

### Assessing gene positions in metabolic networks

To infer which position in a metabolic network a gained gene was likely to occupy, we first mapped each of the 3,022 unique metabolic genes in our study to known metabolic routes in the KEGG MODULE database. Each route in this database is a sequence of steps in the metabolism of a key biomass component; here, the first few steps are catabolic, and the next several steps are anabolic. To each gene, we assigned a position, as follows: (1) first reaction, if the gene corresponded to the first reaction in the route, (2) second reaction, if the gene corresponded to the second reaction, (3) intermediate reaction, for all other reactions (except the last) in the route, and (4) biomass synthesis, for the final reaction in the route. We assumed that genes in categories (1) and (2) were catabolic, and (3) and (4) were anabolic. This assumption is consistent with previous analyses of metabolic gene position [25]. To avoid ambiguity in this analysis, we took two steps. First, we only considered genes which were unique to one route. We verified that relaxing this constraint did not significantly affect our results (supplementary figure 3, where genes present in multiple routes are assigned their most frequently observed position). Second, we excluded short routes (≤ 3 reactions, or steps) from our analysis, since it would be difficult to distinguish catabolism from anabolism in them. We calculated the distribution of gained genes in metabolic networks. For this, on each phylogenetic branch, we calculated the fraction of genes gained along that branch at each metabolic network position. We then averaged the fraction of gained genes at each position across all branches (figure 1c; green bars). As a control, we plotted the expected fraction of gained genes at each position by calculating the average fractions if the genes gained along a branch were a random set of genes, picked from the 3,022 genes in our study; in choosing such random sets, we preserved the number of genes gained along each branch. The average fractions at each position across all branches are plotted as black bars on figure 1c.

### Classifying metabolites as nutrients, byproducts, and biomass components

Of the 8,755 unique metabolites in our study, we classified certain metabolites as nutrients, byproducts and biomass components based on how likely functional roles in metabolic networks. First, to classify between nutrients and byproducts, we curated metabolites based on previously published large-scale metabolic network analyses [37–39]. These analyses used both manual curation and metabolic modeling to distinguish between metabolites that were most likely to be environmentally available nutrients, from those likely to be the metabolic byproducts of other microbes. We found that metabolites on the exterior of metabolic networks were more likely to be nutrients, while those in the interior, byproducts (46 nutrients, 65 byproducts; supplementary table 2). Second, to classify metabolites as biomass components, we used a database of experimentally-verified metabolic models, BiGG [40]. We chose all metabolites listed in the biomass composition of different microbes as biomass components (total 137 metabolites, supplementary table 2).

### Calculating catabolic routes enabled by HGT

We calculated the number of new catabolic routes enabled by HGT along each phylogenetic branch. For this, we first calculated the number of routes in each ancestral metabolic network. We distinguished between routes starting from nutrients (nutrient-driven routes) and byproducts (byproduct-driven routes). We calculated the total number of unique paths in each network that started from nutrients and ended at one of the biomass components; similarly, we calculated the number of paths from byproducts. We used standard network analysis algorithms for these calculations. We then calculated the number of nutrient-driven and byproduct-driven routes in each descendant’s metabolic network. Along each branch, we calculated the difference between the number of nutrient-driven and byproduct-driven routes between the descendant and ancestor. We plotted the distribution of this difference (the number of newly accessible routes) across all branches in figure 2a (nutrient-driven in blue, byproduct-driven in red). To compare the path lengths (number of reaction steps) and energy yields (net number of ATP molecules produced) of the new routes, we did the following along each branch: (1) for path lengths, we compared the lengths of the shortest ancestral path with the shortest new path in the descendant, and asked if a new path was shorter, longer, or of equal length (figure 2b); (2) for energy yields, we compared the net number of ATP molecules produced per nutrient or byproduct, along each route; here also we compared the most ATP-yielding ancestral path with the most ATP-yielding new path, and asked if the new path had a higher, lower or equal yield (figure 2c).

### Detecting phylogenetic events consistent with CGLs and pure gene loss

Along each phylogenetic branch, we asked if there were at least one set of gene gains and losses consistent with coupled gains and losses (CGL-consistent transitions; described in figure 3a) and at least one set consistent with pure gene loss (described in figure 3b). We first calculated all routes that were lost and gained in the descendant (compared with the ancestor) as described the previous section. A new dependency arises when a biomass component can no longer be produced using only the environmentally-available nutrients. We considered the possibility of a dependency for a biomass component, one component at a time. We assumed there was a CGL-consistent transition on a branch if, for any biomass component: (1) the ancestor had only one nutrient-driven route to produce it and zero byproduct-driven routes, (2) the ancestor gained at least one byproduct-driven route to produce it, i.e., there was at least one such route in the descendant, and (3) the ancestor lost the nutrient-driven route during the transition to descendant. We assumed there was a pure gene loss-consistent transition on a branch if, for any biomass component: (1) the ancestor had only one nutrient-driven route to produce it and zero byproduct-driven routes, (2) the ancestor lost this route, and did not gain any byproduct-driven routes, i.e., the descendant had no routes to produce the biomass component. We calculated the fraction of branches where we detected CGL-consistent (figure 4a; green) and pure gene loss-consistent transitions (figure 4a; red). As controls, we calculated the expected fraction of branches with either transitions by using a random set of gains and losses instead (figure 4a; grey bars); in choosing such random sets, we preserved the number of gained and lost genes along each branch.

### Modelling the likelihood of dependency in simulated bacterial communities

Since environmental and community context is crucial to determining whether a given set of gene gains and losses will result in a metabolic dependency, we tested in how many environments and bacterial communities, the observed CGL and pure gene loss events on different branches (identified in the previous section) would result in an actual dependency. For this, we used metabolic models in ∼ 1, 000, 000 simulated environment-community combinations. We chose 1,035 environments, each with 2 of the 46 nutrients in supplementary table 2. We chose 900 communities, with 100 at each level of diversity (from 2 species to 10 species, in steps of 1); each community was a set of bacterial species chosen randomly from the 835 in our study. For each environment-community combination, we calculated the set of byproducts generated by the community by computing which metabolic pathway intermediates each species in the community could produce from the nutrients provided in the environment; we determined this using a popular “scope expansion” algorithm [16, 41].

For each phylogenetic branch, we then asked: in what fraction of environment-community combinations would the descendant evolve a dependency that the ancestor did not have, and through which mechanism — CGLs or pure gene loss? For every level of community diversity, we plotted the fraction of examined cases where we detected a possible dependency through CGLs (figure 4b; green); concurrently we plotted the fraction of cases where the dependency was through pure gene loss (figure 4b; red). In each environment-community combination, we assumed we detected a CGL-mediated dependency if the following conditions were satisfied between the ancestor and descendant for any one biomass component: (1) the ancestor had only one nutrient-driven route to produce it and zero byproduct-driven routes, (2) the nutrient was available in that environment, (3) the ancestor gained at least one byproduct-driven route to produce it, i.e., there was at least one such route in the descendant, (4) the byproduct was available as a community byproduct, and (5) the ancestor lost the coupled nutrient-driven route during the transition to descendant. Similarly, in each environment-community combination, we assumed we detected a pure gene-loss mediated dependency if, for any biomass component: (1) the ancestor had only one nutrient-driven route to produce it and zero byproduct-driven routes, (2) the biomass component was available as a community byproduct, and (3) the ancestor lost the nutrient-driven route during the transition to descendant.

## Data and code availability

All computer code and extracted data files are available at: https://github.com/eltanin4/black_queen_critique.

## Acknowledgements

Part of this work was done at, and supported by, the National Centre for Biological Sciences (NCBS-TIFR), Bengaluru, India. I am grateful to Rohini Subrahmanyam, Deepa Agashe and Saurabh Mahajan for valuable and encouraging discussions. A.G. is supported by the Gordon and Betty Moore Foundation as a Physics of Living Systems Fellow through grant number GBMF4513.

## Author contributions

A.G. conceptualized and designed the research, performed it, analyzed the data, and wrote the paper.

## Supplementary Figures

**FIG. S1.**
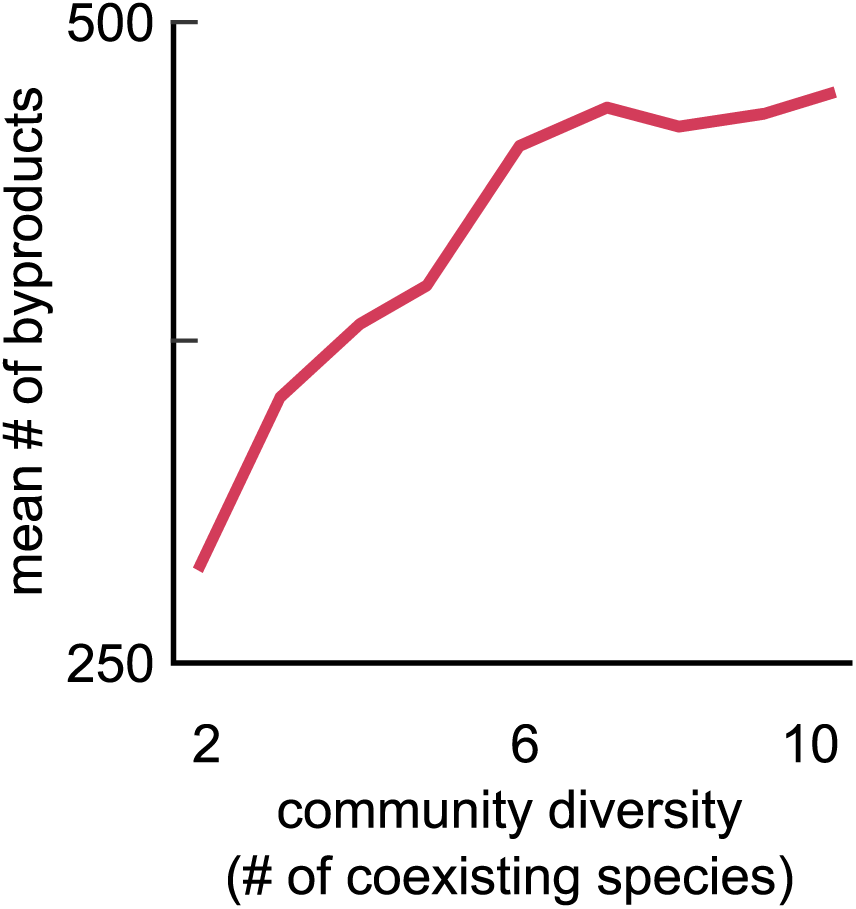
More diverse communities have more community byproducts. Line plot showing the average number of community byproducts in our ∼ 100, 000 environment-community metabolic simulations, as a function of the community diversity. The community diversity is measured by the number of coexisting bacterial species. The community byproducts are calculated using a metabolic network algorithm [41] (see Methods); the average shown is over all environments and communities at a given level of diversity.

**FIG. S2.**
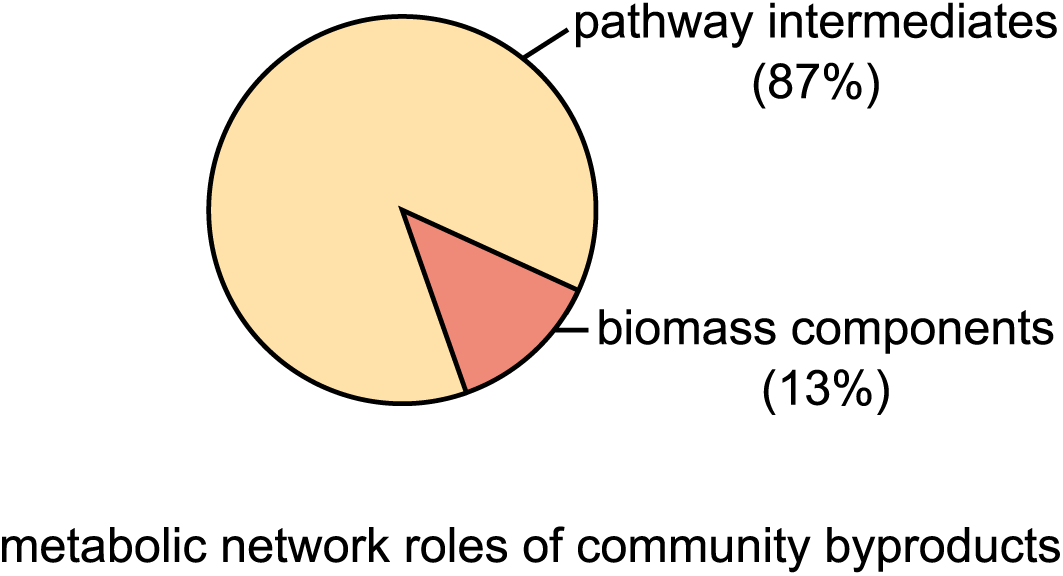
Community byproducts are more likely to be pathway intermediates than end-products (biomass components). Pie chart showing the fraction of all community byproducts in our ∼ 100, 000 environment-community metabolic simulations that were pathway intermediates and pathway end-products (biomass components). Pathway intermediates are more likely to lead to dependencies via coupled gains and losses, because they need to be further metabolized into biomass components by a dependent bacterial species. In contrast, end-products are already biomass components, and do not need to be further metabolized; this makes them more likely to lead to dependencies via pure gene loss.

**FIG. S3.**
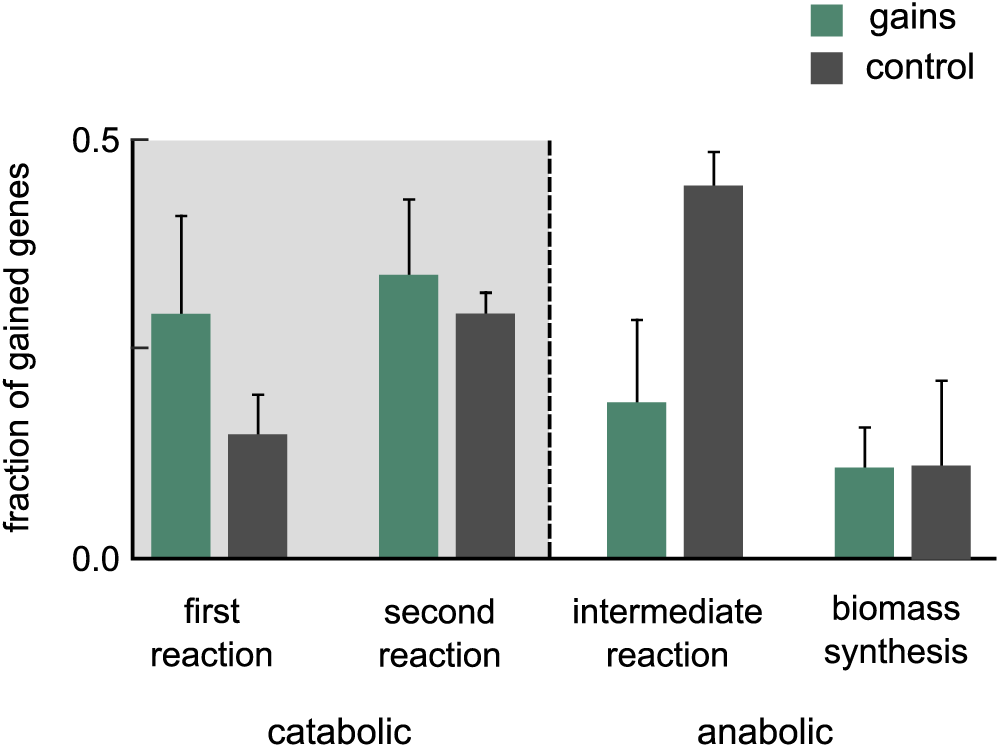
Gained genes are still primarily catabolic, even when network positions are assigned differently to metabolic genes. Bar chart showing the position of genes gained by horizontal gene transfer (HGT) in bacterial metabolic networks, along all 1,669 phylogenetic branches; this chart is similar to figure 1c, except that instead of excluding those genes that participate in multiple metabolic routes, we include them by considering their most frequent metabolic position (catabolic and anabolic). We still exclude those genes that equally occupy catabolic and anabolic positions in this analysis. This alternate assignment choice does not significantly impact our results in figure 1c.

